# An endogenous lentivirus in the germline of a rodent

**DOI:** 10.1101/2022.09.01.506243

**Authors:** Roziah Kambol, Anna Gatseva, Robert J. Gifford

## Abstract

Lentiviruses (genus *Lentivirus*) are complex retroviruses that infect a broad range of mammals, including humans. Unlike many other retrovirus genera, lentiviruses have only rarely been incorporated into the mammalian germline. However, a small number of endogenous retrovirus (ERV) lineages have been identified, and these rare genomic “fossils” can provide crucial insights into the long-term history of lentivirus evolution. Here, we describe a previously unreported endogenous lentivirus lineage in the genome of the South African springhare (*Pedetes capensis*), demonstrating that the host range of lentiviruses has historically extended to rodents (order Rodentia). Furthermore, through comparative and phylogenetic analysis of lentivirus and ERV genomes, considering the biogeographic and ecological characteristics of host species, we reveal broader insights into the long-term evolutionary history of the genus.

## INTRODUCTION

The lentiviruses (genus *Lentivirus*) are an unusual group of retroviruses (family *Retroviridae*) that infect mammals and are associated with a range of slow, progressive diseases in their respective host species groups [1] (**Table 1**). They are most familiar as the genus of retroviruses that includes human immunodeficiency virus type 1 (HIV-1), but the group also includes viruses that infect a broad range of other mammalian groups. Lentiviruses are distinguished from other retroviruses by several characteristic features, including several unique accessory genes, a characteristic nucleotide composition [2, 3], and the capacity to infect non-dividing target cells [4].

**Table 1.**
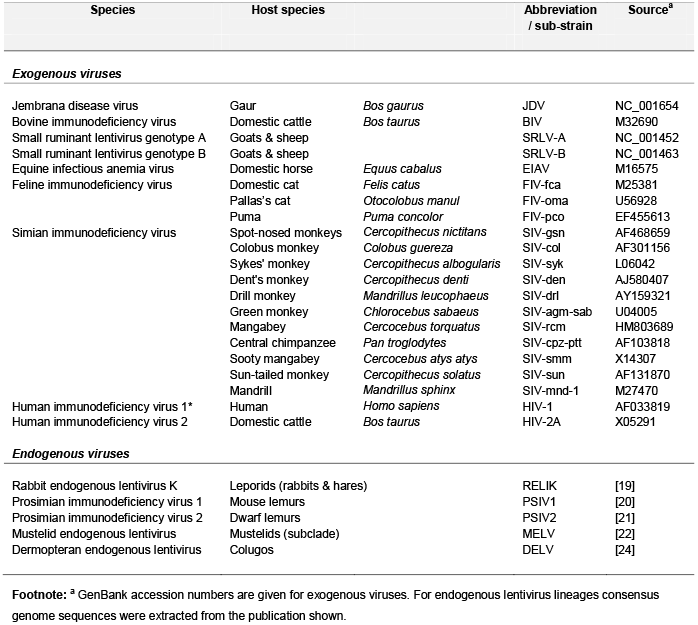
Reference genome sequences of representative lentivirus species.

All retroviruses replicate via an obligate step in which a DNA copy of the viral genome is integrated into a host cell chromosome [5]. The integrated viral genome is flanked at either side by identical long terminal repeat (LTR) sequences (a form referred to as a ‘provirus’), each composed of functionally distinct U3, R and U5 regions. Occasionally, germline cells may be infected and subsequently go on to form viable progeny, so that integrated retroviral proviruses are vertically inherited as host alleles [6]. Such endogenous retroviruses (ERV) insertions are relatively common features in vertebrate genomes [7, 8]. Phylogenetic studies indicate that, following genome invasion, ERVs can increase their germline copy number through a variety of mechanisms, including active replication [9]. However, reflecting their ancient origins, most ERV insertions are genetically fixed and highly degraded by germline mutation. Furthermore, deletion of the entire internal region occurs frequently via homologous recombination between proviral LTRs, leaving behind a single LTR sequence or ‘solo LTR’ [10]. Despite being highly degraded, however, ERVs provide a useful source of retrospective information about the long-term evolutionary interactions between retroviruses and their hosts [11]. For example, identification of orthologous ERV insertions in related species provides a robust means of deriving minimum age calibrations for retrovirus groups, based on host species divergence estimates (which are in part informed by the fossil record) [12]. More broadly, ERV sequences can be used to explore the long-term evolutionary history of ancient - presumably extinct - retrovirus groups [13, 14], and to inform our understanding of their interactions with host genes [15]. ERV sequences can even be used to guide the reconstitution of ancient retrovirus proteins so that their biological properties may be empirically investigated *in vitro* [16–18].

Lentiviruses have only rarely been incorporated into the germline of host species. However, a handful of Lentivirus-derived ERV lineages have now been identified (**Table 1**), and these sequences demonstrate that viruses clearly recognisable as lentiviruses circulated in mammals many millions of years ago. For example, rabbit endogenous lentivirus K (RELIK) insertions were found to occur at orthologous positions in the rabbit (*Oryctolagus cuniculus*) and hare (*Lepus europaeus*) genomes, demonstrating that genome invasion occurred prior to divergence of these species ~12 million years ago (Mya) [12, 19]. Endogenous lentiviruses have also been identified in lemurs (family Lemuridae) [20, 21]; mustelids (family Mustelidae) [22, 23]; and dermopterans (order Dermoptera - a group of arboreal gliding mammals native to Southeast Asia) [24–26]. Together, these sequences provide a range of minimum age calibrations in the Miocene epoch (23.5-5.3 Mya), based on host species divergence date estimates derived from the fossil record [11, 22, 25]. Widespread circulation among mammals is further supported by estimates derived via application of a molecular clock, some of which extend into the Eocene epoch (56-33.9 Mya) [24, 26].

In this study we perform comprehensive screening of published mammalian genomes and identify a previously unreported endogenous lentivirus lineage in the genome of the South African springhare (*Pedetes capensis*), demonstrating that lentivirus host range extends to rodents. Furthermore, through comparative and phylogenetic analysis, incorporating all available data, we provide broader insight into the origins and long-term evolutionary history of lentiviruses.

## MATERIALS & METHODS

### Genome screening in silico

We used database-integrated genome screening (DIGS) [27] to derive a non- redundant database of lentivirus-derived ERV loci contained in published genome sequence assemblies. In DIGS, the output of systematic, sequence similarity search-based ‘screens’ is captured in a relational database. The DIGS tool [27] is a Perl-based framework in which the Basic Local Alignment Search Tool (BLAST) program suite (version 2.2.31+) [28] is used to perform systematic similarity searches of sequence databases (e.g., genome assemblies) and the MySQL relational database management system (MySQL Community Server version 8.0.30) is used to record and organise output data. WGS data of 431 mammalian species were obtained from the National Center for Biotechnology Information (NCBI) genome database [29] (**Table S1**). Query polypeptide sequences were derived from representative lentivirus species (**Table 1**). DNA sequences in WGS assemblies that disclosed significant similarity to lentivirus queries (as determined by BLAST *e-*value) were classified via comparison to published retrovirus genome sequences (again using BLAST). Consensus genome sequences for endogenous lentivirus lineages were extracted from the supplementary material of associated publications, as follows: RELIK [19]; PSIV1 [20]; PSIV2 [21]; MELV [22]; DELV [24].

We compiled a set of endogenous lentivirus loci (**Table S2**) by using structured query language) to filter screening the classified, non-redundant results of >130,000 searches, selecting matches based on their degree of similarity to lentivirus reference sequences, or the taxonomic characteristics of the species in which they occur. Using this approach we separated putatively novel lentivirus ERV loci from both (i) orthologs or paralogs of previously characterised lentivirus ERVs, and (ii) non-lentiviral sequences that cross-matched to lentivirus probes due to shared ancestry (e.g., clade II ERVs) [30, 31]. We confirmed that putative novel lentivirus ERVs were indeed derived from lentiviruses (rather than other, related retroviruses) through phylogenetic and genomic analysis as described below.

### Phylogenetic and genomic analysis

Nucleotide and protein phylogenies were reconstructed using maximum likelihood (ML) as implemented in RAxML (version 8.2.12) [32]. Protein substitution models were selected via hierarchical maximum likelihood ratio test using the PROTAUTOGAMMA option in RAxML. To estimate the ages of solo LTRs we measured divergence from an LTR consensus sequence and applied a neutral rate calibration, as described by Subramanian *et al*. [33]. We used Se-Al (version 2.0) to visualise alignments and create consensus sequences [34].

## RESULTS & DISCUSSION

We systematically screened WGS data representing 431 mammalian species (**Table S1**) for endogenous lentivirus loci using similarity search-based approaches We identified a total of 842 distinct lentivirus-derived ERV loci, most of which represented members of previously described lentivirus ERV lineages (**Table 2,** [35]). However, we also identified lentivirus-derived sequences in the genome of a species group in which they have not previously been described – rodents (order Rodentia).

**Table 2.**
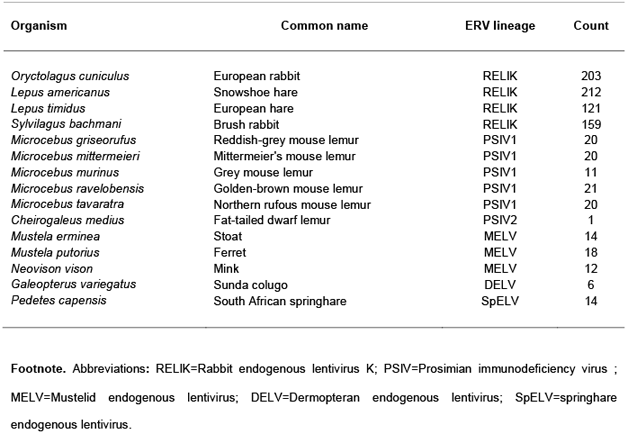
Endogenous lentivirus loci detected via screening.

Matches to lentiviral Gag and Pol proteins were identified in WGS data of the South African springhare (*Pedetes capensis*), and phylogenetic analysis of the reverse transcriptase (RT) coding region encoded by these ERVs demonstrates that they group within the diversity of previously described lentivirus species (**Fig. S1a**). Initially, only four copies of Springhare endogenous lentivirus (SpELV) were identified in the *P. capensis* genome. However, we were able to identify the 5’ LTR of a partial provirus sequence by using upstream flanking sequence as a query in BLASTn-based searches of the *P. capensis* genome assembly. This revealed the presence of a repetitive sequence showing the characteristic features of a retroviral LTR (i.e., ~500 nucleotides in length with terminal TG and CA dinucleotides) in the expected position upstream of the Gag ORF. Using this LTR sequence as input for screening enabled us to identify another 10 SpELV loci represented by solo LTR sequences (**Table 3)**. We generated a consensus SpELV genome using all fourteen loci identified in our screen (**Fig. S2**). We did not identify an envelope (*env*) gene associated with any SpELV insertions, nor did we identify any contigs containing complete proviruses with paired LTR sequences. Furthermore, because the longest provirus sequence we identified was truncated in *pol* we could not determine whether any accessory genes might have been encoded downstream of this gene. Nonetheless, the partial genome obtained in our analysis exhibits the characteristic features of lentivirus genomes, including (i) a primer-binding site specific for tRNA Lysine (**Fig. S3**); (ii) a Pro-Pol ORF expressed via - 1 ribosomal frameshifting (**Fig. S3**); (iii) an adenine-rich (34%) genome (**Fig. S4**) containing few CpG dinucleotides (0.29%); (iv) a putative *trans*-activator response (TAR) element (**Fig. S2, Fig. S3**). We estimated the age of the SpELV lineage utilising a molecular clock-based approach in which divergence is calculated by comparing individual LTR sequences to an LTR consensus [33]. We obtained age estimates in the range of 8-18 Mya for SpELV loci (**Table 3**), consistent with an origin in the Middle Miocene.

**Table 3.** Springhare endogenous lentivirus loci.

| ERV locus<sup>a</sup> | Structure | Age (Mya)<sup>b</sup> | Contig | Orientation | Start | End |
| --- | --- | --- | --- | --- | --- | --- |
| SpELV.1-PedCap | LTR-Gag-Pol | 8,750,000 | VMDO01011022.1 | + | 63971 | 68252 |
| SpELV.2-PedCap | LTR-Gag | 12,500,000 | VMDO01050306.1 | + | 2112 | 3978 |
| SpELV.3-PedCap | Gag | ND | VMDO01082106.1 | - | 284 | 997 |
| SpELV.4-PedCap | Gag | ND | VMDO01088624.1 | - | 445 | 1155 |
| SpELV.6-PedCap | LTR | 10,084,925 | VMDO01001729.1 | - | 65134 | 65652 |
| SpELV.14-PedCap | LTR | 11,194,029 | VMDO01006229.1 | + | 41369 | 41857 |
| SpELV.15-PedCap | LTR | 11,460,554 | VMDO01001857.1 | + | 174371 | 174854 |
| SpELV.16-PedCap | LTR | 11,460,554 | VMDO01001488.1 | + | 108014 | 108496 |
| SpELV.17-PedCap | LTR | 8,528,784 | VMDO01000902.1 | + | 172296 | 172776 |
| SpELV.18-PedCap | LTR | 11,143,410 | VMDO01003412.1 | + | 3534 | 4013 |
| SpELV.21-PedCap | LTR | 9,594,882 | VMDO01035581.1 | - | 11304 | 11783 |
| SpELV.24-PedCap | LTR | 11,388,286 | VMDO01046849.1 | + | 1434 | 1906 |
| SpELV.25-PedCap | LTR | 13,592,750 | VMDO01015080.1 | + | 21755 | 22226 |
| SpELV.28-PedCap | LTR | 17,665,952 | VMDO01001033.1 | - | 148080 | 148542 |

**Footnote:** <sup>a</sup> Loci were assigned unique identifiers (IDs) following a standard nomenclature system [45]. PedCap=*Pedetes capensis*. The ages of LTR-encoding elements was estimated by measuring divergence from an LTR consensus sequence and applying a neutral rate calibration, as described by Subramanian *et al.* [33].
ND=not done. SpELV=Springhare endogenous lentivirus.

We used maximum likelihood-based phylogenetic approaches to reconstruct the evolutionary relationships between contemporary lentiviruses and the extinct lentiviruses represented by ERVs. Phylogenetic trees clearly separate the Lentiviruses into two robustly supported subclades (**Fig. 1**). One (here labelled ‘Archaeolentivirus’) contains SpELV together with dermopteran endogenous lentivirus (DELV) which occurs in the germline of colugos [24–26]. A second (here labelled ‘Neolentivirus’) contains all other endogenous lentivirus lineages and all known contemporary lentiviruses. We obtained relatively high support for internal branching relationships within the Neolentivirus clade – reconstructions support the existence of a distinct ‘primate’ group of neolentiviruses containing both simian and prosimian sub-lineages, and an ‘artiodactyl’ group incorporating both the bovine lentiviruses and the small ruminant lentiviruses. In addition, the primate lentiviruses group separately from all other neolentiviruses, which together constitute a ‘grasslands-associated’ clade comprised of lentiviruses that infect(ed) grassland-adapted host species.

**Figure 1.**
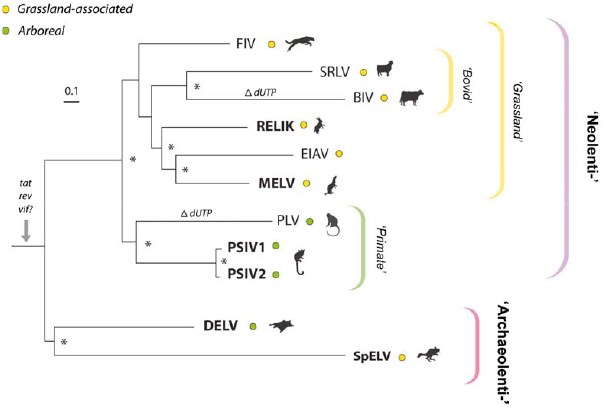
Phylogenetic relationships within the Lentivirus genus. Maximum likelihood phylogeny showing reconstructed evolutionary relationships between all known lentivirus species, including the extinct species represented by endogenous lentiviruses. Brackets to the left indicate proposed subgroupings within genus *Lentivirus*. Coloured circles adjacent virus taxa labels indicate the ecological characteristics of the associated host species (grassland-dwelling or arboreal) as shown in the key top right. The phylogeny is midpoint rooted for display purposes and was reconstructed using a multiple sequence alignment spanning 1405 amino acid residues of the Gag-Pol polyprotein and the RT-Rev substitution model [46]. The scale bar shows evolutionary distance in substitutions per site. Asterisks indicate nodes with bootstrap support >70% (1000 replicates). **Abbreviations**: DELV=Dermopteran endogenous lentivirus; SpELV=springhare endogenous lentivirus. RELIK=Rabbit endogenous lentivirus type K; Mustelidae endogenous lentivirus (MELV); BIV=Bovine immunodeficiency virus; SIV=Simian immunodeficiency virus; EIAV=equine infectious anaemia virus; FIV=Feline immunodeficiency virus; Human immunodeficiency virus=HIV; SRLV=small ruminant lentivirus; PSIV=Prosimian immunodeficiency virus.

To examine the distribution and diversity of lentiviruses in the context of host evolution, we plotted information related to (i) lentivirus distribution and (ii) host biogeographic range onto a time-calibrated phylogeny of boreoeutherian hosts (**Fig. 2**). This revealed that age estimates obtained from the genomic fossil record (either through the identification of ancient orthologs, or via the application of a molecular clock) are consistent with other calibrations in deep time that can be tentatively inferred from ancestral biogeographic distributions by parsimoniously assuming limited transfer of virus between major biogeographic regions and distantly related host groups. Lentiviruses are known to cross species barriers quite frequently [36, 37], but transmission between large phylogenetic distances (e.g., distinct taxonomic orders of mammals) has never been reported and is unlikely to be common based on current understanding of the barriers to zoonotic transfer [38]. Evidence from orthology and molecular clock-based analyses supports the presence of DELV in Asia (the only region where colugos occur) up to 60 Mya – i.e., throughout most of the Cenozoic Era [26]. identification of a DELV-related virus in springhares – which evolved in the African subcontinent – implies the presence of archaeolentiviruses in ancestral mammals >80 Mya [39]. Notably, other groupings within the Neolentivirus subclade are also consistent with late Cretaceous origins predating the subordinal divergences of major placental mammal groups. For example, the existence of an ancient primate lineage, incorporating both lemurs, apes, and monkeys (**Fig. 1**) is consistent with a parsimonious scenario under which lentiviruses were present in the common ancestor of all primates and arrived in Madagascar with founder populations of ancestral lemurs ~60 Mya [40, 41].

**Figure 2.**
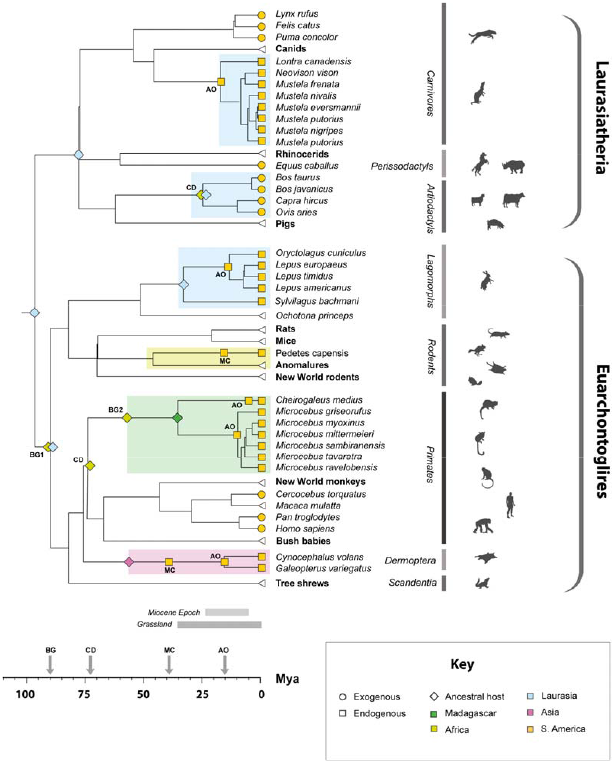
An updated timeline of lentivirus evolution. A time-calibrated phylogeny of mammalian species showing the known extent of association between lentiviruses and mammals, based on data obtained from TimeTree [47]. The scale bar shows time in millions of years before present. Brackets and bars to the right of taxa labels indicate host taxonomic groups. Coloured squares on terminal nodes indicate that host species associated with endogenous lentiviruses (squares) or exogenous lentiviruses (circles). The timeframe of endogenous lentivirus presence in each mammalian lineage is indicated by shaded boxes underneath clades, with colours indicating biogeographic associations of hosts within each clade following the key. White triangles at tree tips indicate host species or groups that have not yet been associated with any lentiviruses (endogenous or exogenous). Two-letter codes adjacent internal markers indicate the type of calibration being shown, as follows (AO=identification of an ancient ortholog; MC=application of a molecular clock to neutrally diverging sequences; CD=assumption of codivergence with hosts; BG=assumption of presence in biogeographic area inhabited by ancestor of species groups that are now biogeographically separated – note that this assumes no transfer between the respective regions identified by derived host species). Colours on diamond-shaped node markers indicate the known biogeographic range of ancestral hosts, as indicated in the key. The biogeographic range of the springhare-colugo ancestor (BG1) is uncertain (hence two regions are shown). The colonisation of isolated Madagascar by lemurs (BG2) is thought to have occurred ~60 million years ago (Mya) [40, 41].

At the same time, it seems clear that transmission of lentiviruses between phylogenetically distant mammal groups has occurred in the past. For example, the ‘grasslands-associated’ subclade contains viruses and paleoviruses that infect (or infected) phylogenetically distinct host species groups that share grassland habitat. It includes equine infectious anaemia virus (EIAV) which infects horses, and two ERV lineages - RELIK (found in leporids) and MELV (found in mustelids) [42–44]. Notably, the grassland adaptation of these three host species groups took place in a similar time-period (early-to-middle Miocene) in interconnected biogeographic areas (Laurasia and Africa) [42–44] (**Fig. 2**), suggesting that the connections between the viruses in this clade could reflect inter-order transmission events that took place in a shared habitat.

## CONCLUSIONS

We describe a novel endogenous lineage in the genome of the South African springhare. The identification of SpELV demonstrates that lentivirus host range has historically extended to rodents. Through comparative and phylogenetic analysis of modern and ancient lentivirus genomes, we show that the *Lentivirus* genus incorporates at least two major subclades and reveal evidence it originated >80 Mya. In addition, we reveal phylogenetic evidence that transmission of lentiviruses between distantly related mammalian groups (i.e., distinct orders) has historically occurred in shared habitats.

## SUPPLEMENTARY FIGURE LEGENDS

**Figure S1.**
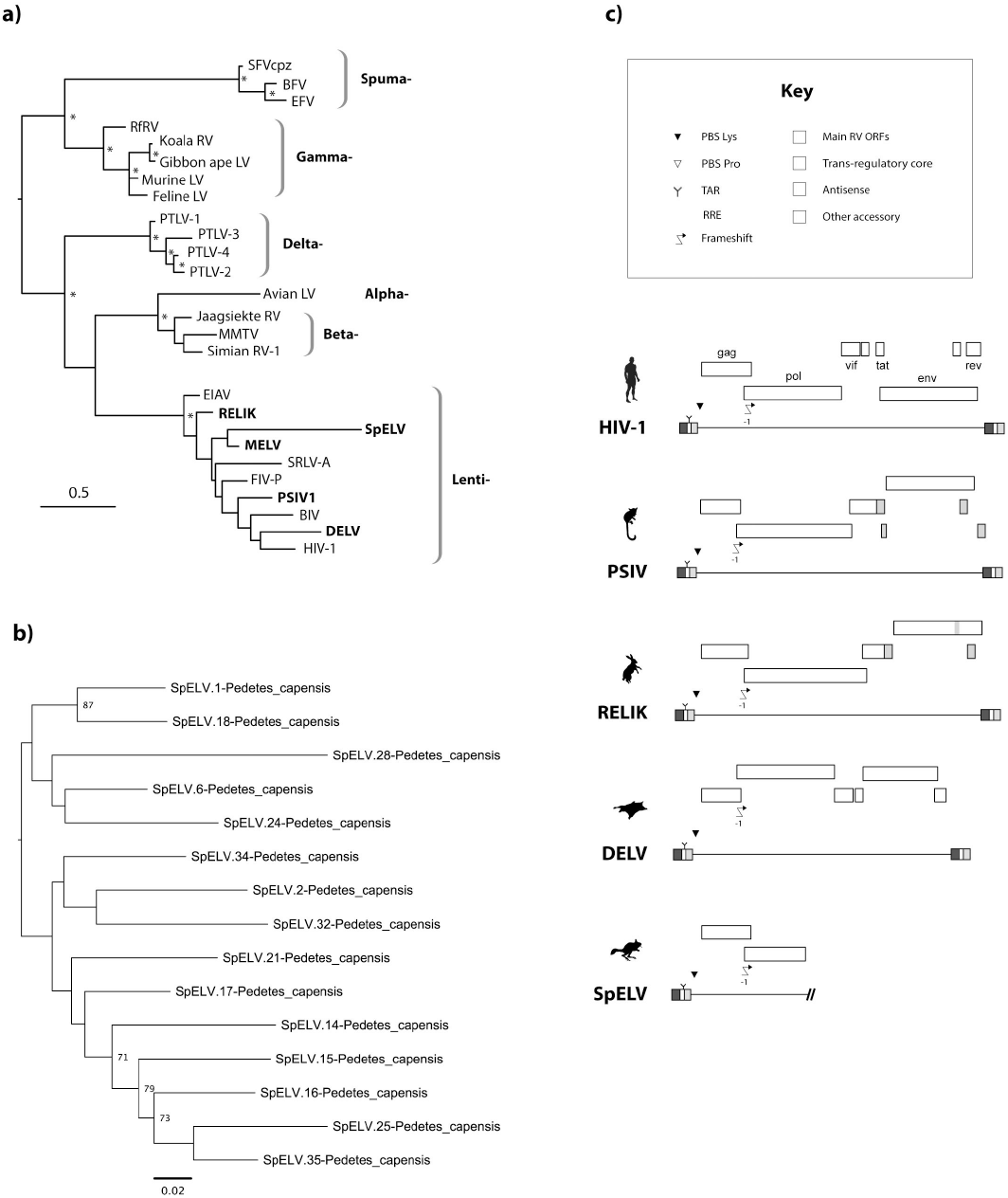
Phylogenetic and genomic characteristics of springhare endogenous lentivirus. **(a)** Maximum likelihood (ML) phylogeny based on an alignment of reverse transcriptase (RT) protein sequences and showing the reconstructed evolutionary relationships between lentiviruses and other retroviruses. Asterisks indicate nodes with bootstrap support >70% (1000 replicates). The scale bar shows evolutionary distance in substitutions per site. **(b)** ML phylogeny showing reconstructed evolutionary relationships between SpELV long terminal repeat (LTR) sequences. Numbers next to nodes indicate bootstrap support values (1000 replicates). The scale bar shows evolutionary distance in substitutions per site. **(c)** Consensus genome structures of ancient lentiviral paleoviruses. **Abbreviations:** DELV=Dermopteran endogenous lentivirus; RELIK=Rabbit endogenous lentivirus type K; Mustelidae endogenous lentivirus (MELV); BIV=Bovine immunodeficiency virus; SIV=Simian immunodeficiency virus; FIV=Feline immunodeficiency virus; Human immunodeficiency virus=HIV; Prosimian immunodeficiency virus=PSIV; RV=Retrovirus; LV=Leukemia virus.

**Figure S2.**
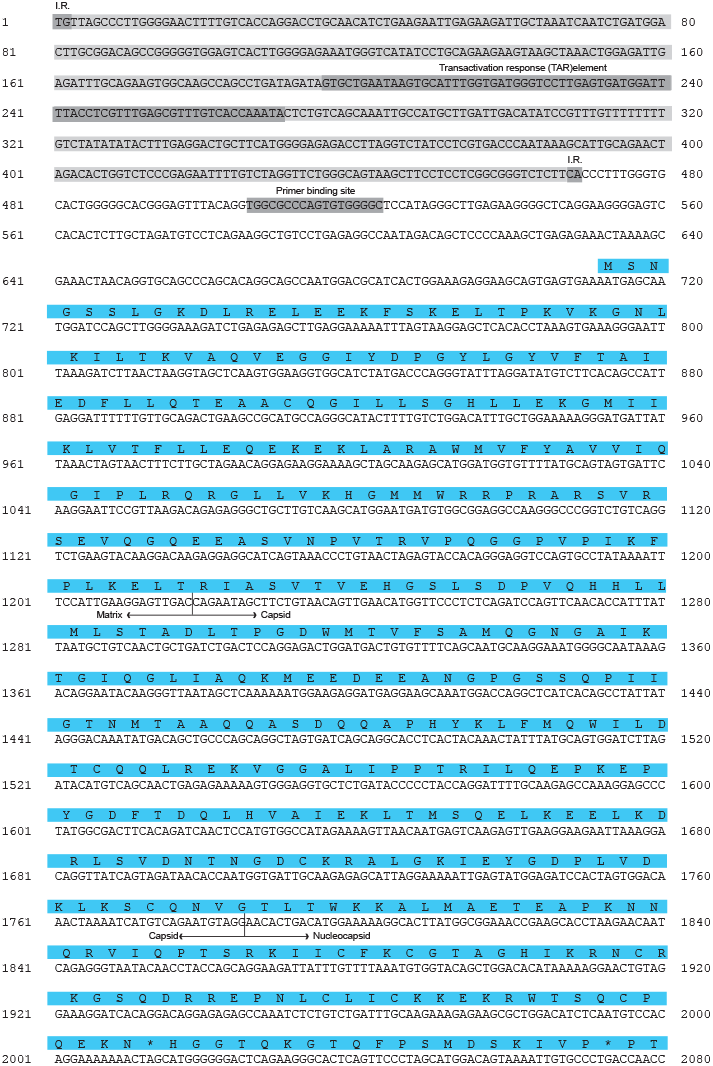
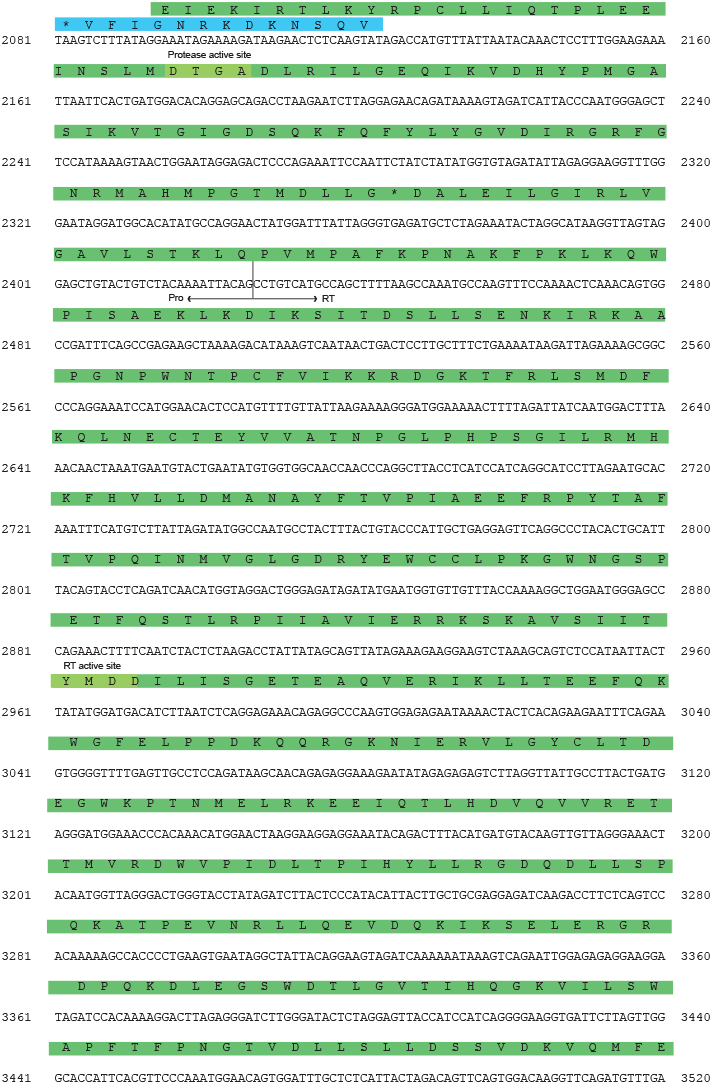
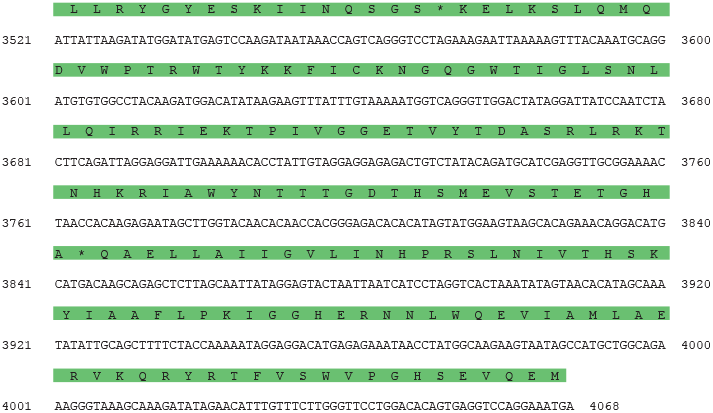
The SpELV consensus sequence. Inverted repeats present at the ends of the 5’ long terminal repeat (LTR) sequence are highlighted in light grey. Regions of nucleic acid secondary structure, the transactivation responsive (TAR) element and primer binding site (PBS) are highlighted in dark grey. The locations of the proteins encoded by the *gag* and *pol* genes were determined by homology to the DELV consensus sequence [24–26].

**Figure S3.**
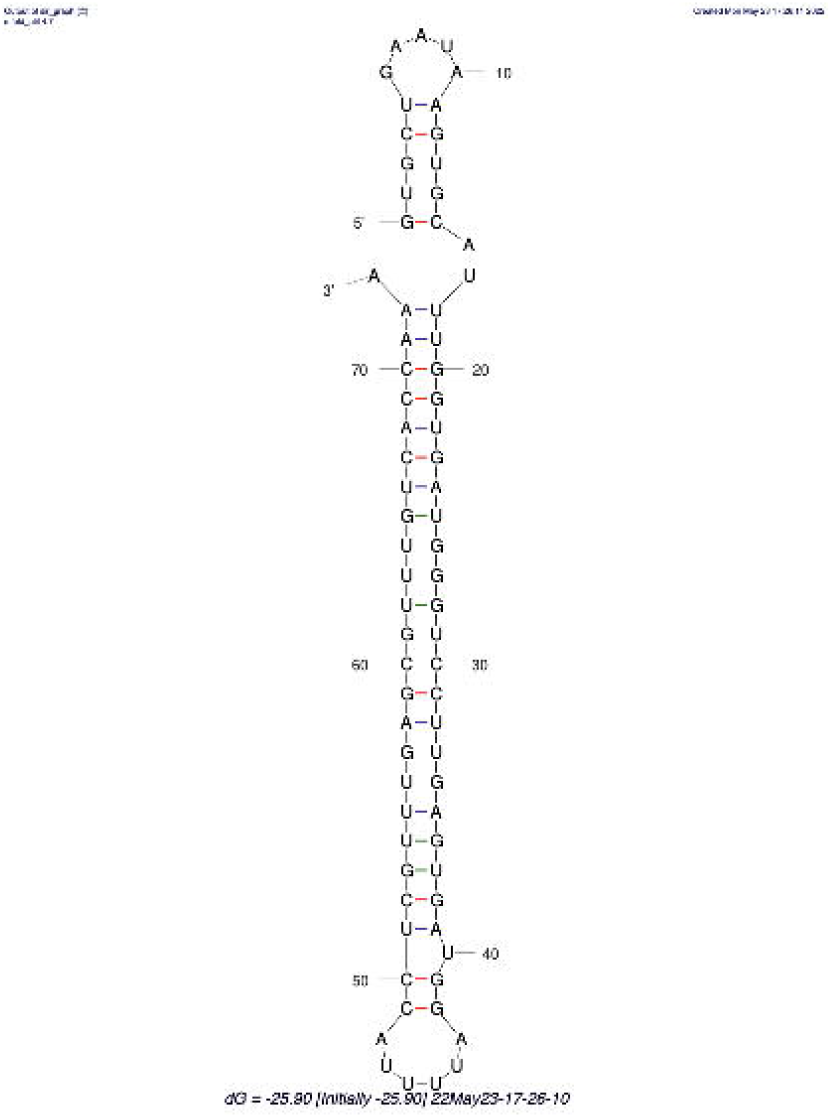
The putative SpELV TAR (transactivation responsive region) element. Secondary structures were predicted using the MFOLD thermodynamic folding algorithm [48] and assessed by comparison to well-characterised examples in other lentiviruses.

**Figure S4.**
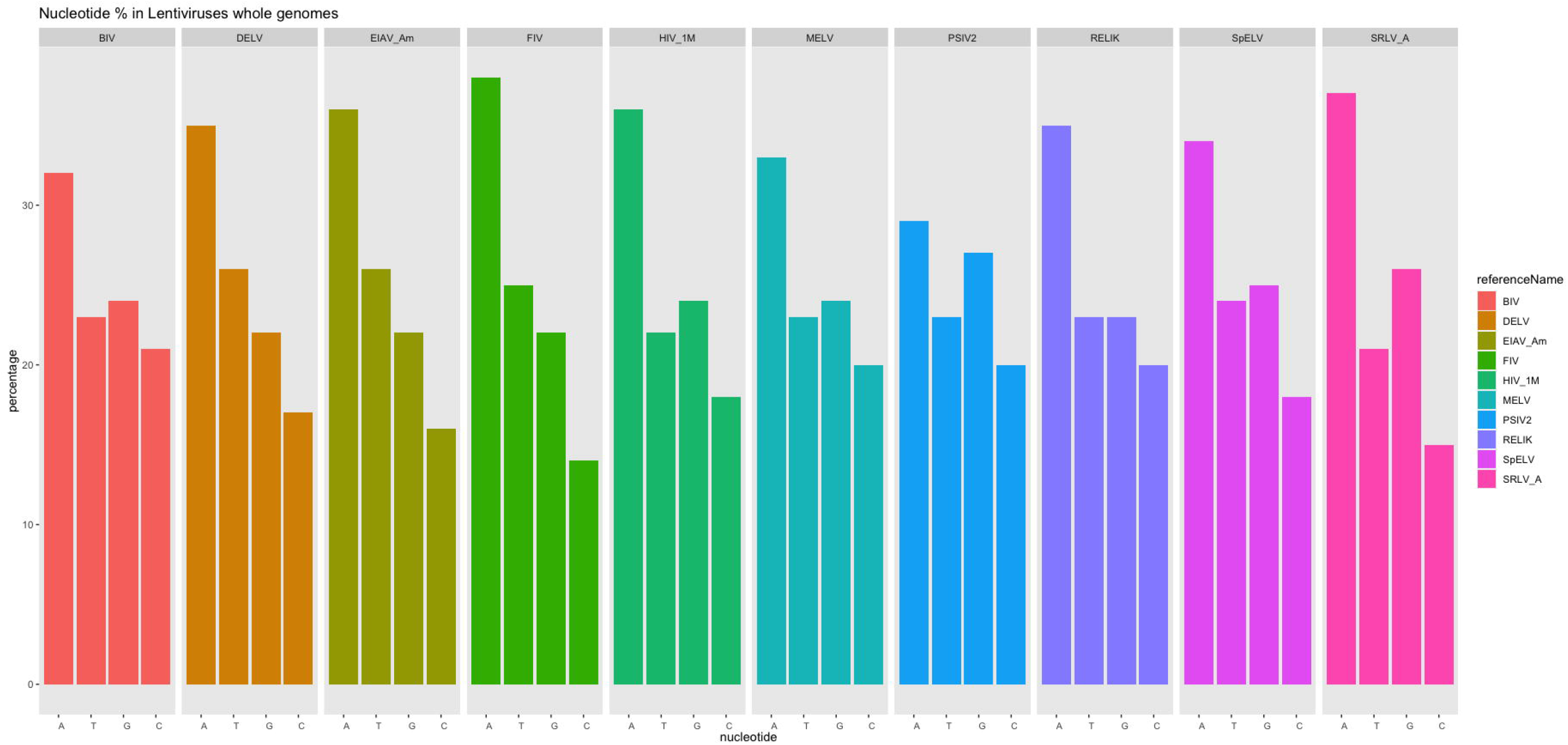
Nucleotide compositional bias in lentivirus genomes. Nucleotide composition of whole genomes of Lentiviruses were normalised to length and plotted as percentages using R in R Studio (version 4.2.1). Reference genome sequences for each virus correspond to those given in **Table 1**. Bovine immunodeficiency virus (BIV), Dermopteran endogenous lentivirus (DELV), Equine infectious anaemia virus American strain (EIAV_Am), Feline immunodeficiency virus (FIV), Human immunodeficiency virus 1 (HIV_1M), Mustelidae endogenous lentivirus (MELV), Prosimian immunodeficiency virus 2 (PSIV); Rabbit endogenous lentivirus type K (RELIK), Springhare endogenous lentivirus (SpELV), Small ruminant lentivirus A (SRLV_A); Adenine (A), Guanine (G), Cytosine (C), Thymine (T).

## DECLARATIONS

### Ethics approval and consent to participate

Not applicable

### Consent for publication

Not applicable

### Availability of data and materials

All data are openly available in the Lentivirus-GLUE project hosted on GitHub:

https://giffordlabcvr.github.io/Lentivirus-GLUE/

## Acknowledgments

We thank Daniel Blanco-Melo, Anne Emory, Ron Swanstrom and Greg Towers for helpful discussions.

## Authors’ contributions

Conceptualization, R.J.G.; methodology and validation, A.G., R.K., and R.J.G.; formal analysis, A.G., R.J.G.; writing—original draft preparation, R.J.G.; writing—review and editing, A.G., R.J.G., and R.K.; visualization, A.G., R.J.G.; supervision, R.J.G.; project administration, R.J.G.; data curation, R.J.G. All authors have read and agreed to the published version of the manuscript.

